# Classification of the lower limb motor imagery and execution using high vs. low channel EEG devices

**DOI:** 10.64898/2026.02.10.704620

**Authors:** Parsa Bahramsari, Saeed Behzadipour

## Abstract

Brain–computer interfaces (BCIs) translate brain signals into commands for external devices, with motor imagery (MI) BCIs decoding imagined movements to aid neurorehabilitation. Although high-channel EEG offers rich data, such systems are bulky and impractical for everyday use. This study assesses whether a low-channel, consumer-grade headset (Muse) can match a clinical-grade system (OpenBCI) in classifying lower limb MI and motor execution (ME). Six healthy volunteers performed left and right knees and ankles MI and ME tasks while EEG was recorded concurrently from both devices. Signals were band-pass filtered (8–30 Hz), segmented into overlapping one second windows, and features were extracted across time, frequency, and time-frequency domains. Feature dimensionality was reduced via mutual information-based minimum redundancy maximum relevance and principal component analysis. Five classifiers (support vector machine, linear discriminant analysis, k nearest neighbors, random forest, and AdaBoost) were applied to nine binary discrimination scenarios and evaluated with 10-fold cross-validation via 100 Monte Carlo iterations. Frequency domain features, particularly those derived from Welch’s power spectral density, were most frequently selected. Mutual information analysis indicated that C3 and C4 electrodes were most informative for OpenBCI, while in Muse, the channels contributed more evenly, except in laterality classification scenarios, where TP9 played a key role. OpenBCI outperformed Muse in classifier-based accuracy with superiority ranging from 0.4% to 4.8%, while task-based differences were more variable, ranging from -0.3% to 8.7%. Despite its lower spatial resolution, the Muse system achieved competitive performance, especially in motor vs. rest tasks, and shows promise as an affordable, user-friendly alternative for home-based neurorehabilitation BCIs.

## 1. Introduction

Brain computer interfaces (BCIs) are systems that measure central nervous system (CNS) activity to control external devices that would replace, restore, enhance, supplement, or improve the natural CNS output [1]. Electroencephalography (EEG) is widely used for BCI applications due to its high temporal resolution, ease of use, and affordability [2]. EEG-based BCIs are designed to monitor brain activity and translate it into control signals that can operate external devices. One prominent category of tasks used in BCIs is motor tasks, which include motor imagery (MI) tasks, where individuals imagine movement without actual execution, and motor execution (ME) tasks, where physical movement is performed. In MI-BCI systems, the brain’s cortical activity associated with imagined movement is captured and translated into control signals for devices, offering potential for rehabilitation and assistive technology in conditions such as stroke, spinal cord injury, or neurodegenerative diseases [2, 3,4]. According to the Hebbian principle, which states that “neurons that fire together wire together”, the repeated engagement of specific neural circuits through MI can strengthen synaptic connections, facilitating neuroplastic changes and assisting in motor recovery [5]. EEG signals can be specifically trained to detect motor tasks, providing a pathway for restoring lost functions. Such systems may also be used to guide brain activity back to its physiological state, helping users regain motor control and improve their functional capabilities [6].

Various studies have explored the use of EEG for classifying lower limb MI tasks emphasizing the role of channel count and spatial distribution in improving classification accuracy. For instance, Zuo et al. used 32 channels to classify left-leg vs. right-leg MI, reporting average accuracies ranging from 51.43% to 86.41% across approaches [7]. Ma et al. used 64 channels to classify four lower-limb classes of left-leg flexion MI, left-leg extension MI, right-leg flexion MI, and right-leg extension MI, and reported an average accuracy of 62.68% [8]. Peng et al. used 64 channels to classify leg-raise MI vs. rest, reporting an average accuracy of 88.43% [9]. While these studies demonstrate promising classification results, the high-channel EEG devices used in these experiments, despite their effectiveness in improving MI-BCI performance, introduce significant practical challenges that limit their applicability in real-world, clinical, or home environments. Setting up these EEG devices can be challenging, as precise electrode placement often requires technical expertise, limiting their usability in everyday or unsupervised settings. Optimal signal quality typically involves conductive gels, skin abrasion, or additional preparations, making the process uncomfortable and time-consuming. Furthermore, wires and cables restrict user movement, especially problematic in motor-task BCIs, and can introduce signal artifacts that reduce EEG recording quality [10, 11, 12]. Although these high-channel EEG devices offer better spatial resolution, potentially leading to improved classification performance, they remain difficult to use, especially in non-clinical settings. These challenges highlight the need for more practical EEG-based BCI solutions that balance classification performance with usability and accessibility for end-users in real-world applications.

In this research, we collected EEG data using two distinct EEG systems: a consumer-grade EEG device designed for usability and real-world applications, and a clinical system allowing greater flexibility in data acquisition. The consumer-grade EEG device, offering minimal setup, high usability, and comfort, making it a practical solution for home-based neurofeedback, brain training, and lightweight BCI interactions. However, its limited channel setup and electrode placement in less motor-relevant regions may reduce its effectiveness for detection of motor tasks. In contrast, clinical system allows customizable electrode placement, enabling data acquisition from any desired location, which enhances its ability to capture motor tasks. However, its use involves complex electrode setup, conductive gels, wired connections, and potential signal instability during movement, making it less convenient for practical use. Both systems were tested under a unified experimental protocol to facilitate direct comparison of their ability to detect and classify lower limb motor tasks. The tasks under investigation included knees MI and ME, as well as ankles MI.

The primary goal of this study was to investigate the feasibility of consumer-grade EEG device to develop a MI-BCI in comparison to a clinical device. In other words, the main question was whether the performance loss due to the lower spatial resolution of the signal is worth the simplicity of application. Additionally, this research assesses the impact of features, channels, and classifiers on classification accuracy. Beyond device comparison, our study contributes to the ongoing development of BCI technology for neurorehabilitation by deepening our understanding of lower limb motor tasks classification.

## 2. Methods

A subject-specific analysis was conducted to compare the two EEG systems in detecting and classifying lower-limb motor tasks. An overview of the method is shown in “Figure 1” and the details are given in the following subsections.

**Figure 1.**
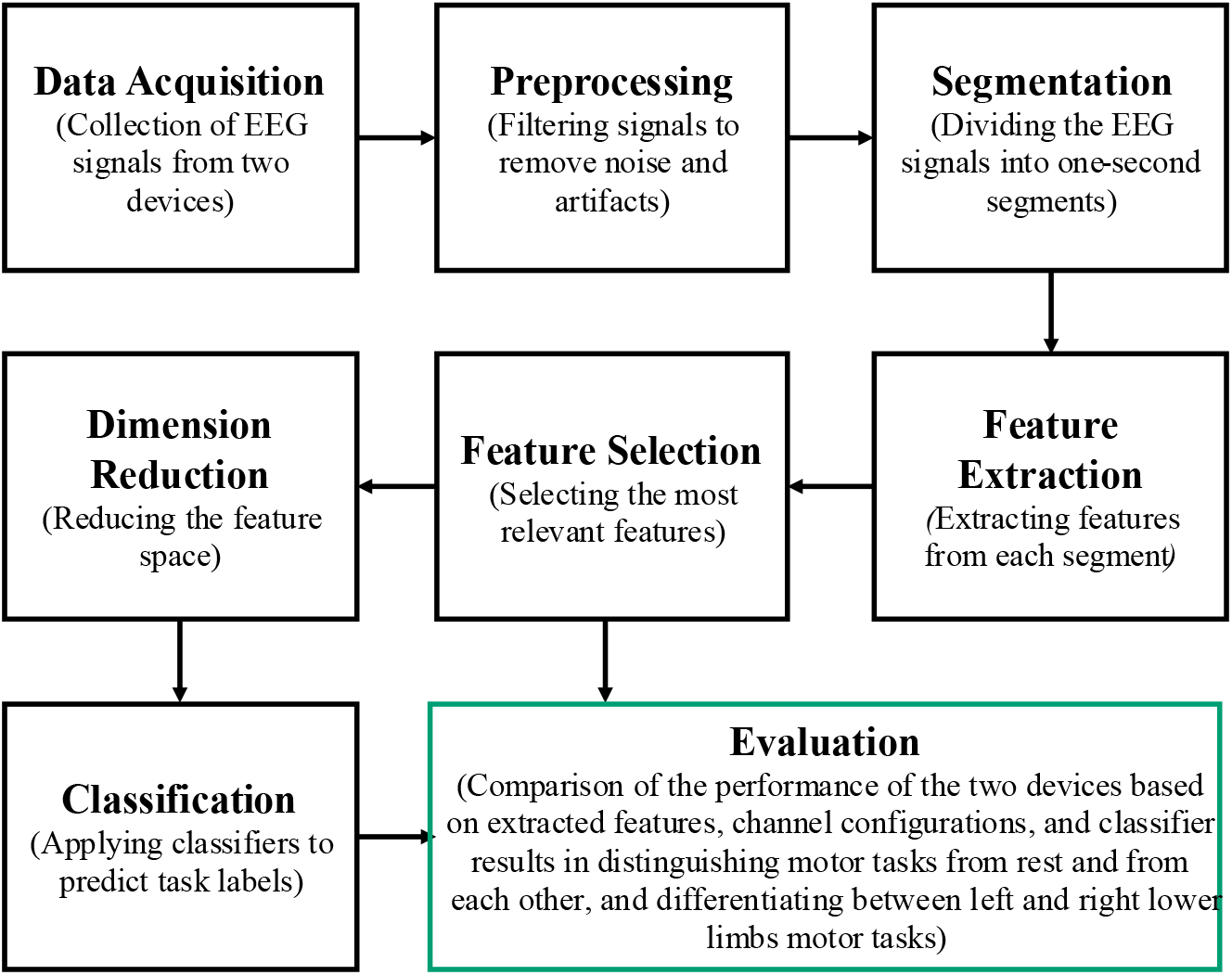
Methodological pipeline for the analysis of EEG signals

### 2.1. EEG Recordings

This study utilized two EEG devices with distinct configurations: a 4-channel Muse headband and an 8-channel OpenBCI system. The Muse headband refers to the Muse 2016 EEG device (InteraXon Inc., Toronto, Ontario, Canada), which records data at 256 Hz using four dry electrodes placed at channels TP9, AF7, AF8, and TP10, based on the modified expanded 10–20 system [13], with FPz used as the reference. Data were transmitted via Bluetooth and captured using the MuseLSL package [14] in Python. The OpenBCI system denotes the full setup comprising the OpenBCI Cyton 8-channel board (OpenBCI Inc., Brooklyn, New York, United States), eight MCScap-E Ag/AgCl sintered electrodes, and an MCScap 10–20 textile cap (Medical Computer Systems Ltd., Zelenograd, Moscow, Russia). It recorded from the channels C1, C3, FC1, CP1, C2, C4, FC2, and CP2, which are closely associated with motor-related cortical areas, particularly the sensorimotor and premotor regions involved in movement planning and execution [15]. Signals were referenced to Cz and sampled at 250 Hz, with data transmitted to a USB dongle and recorded via OpenBCI GUI. “Figure 2” illustrates each device’s setup and channel configuration.

**Figure 2.**
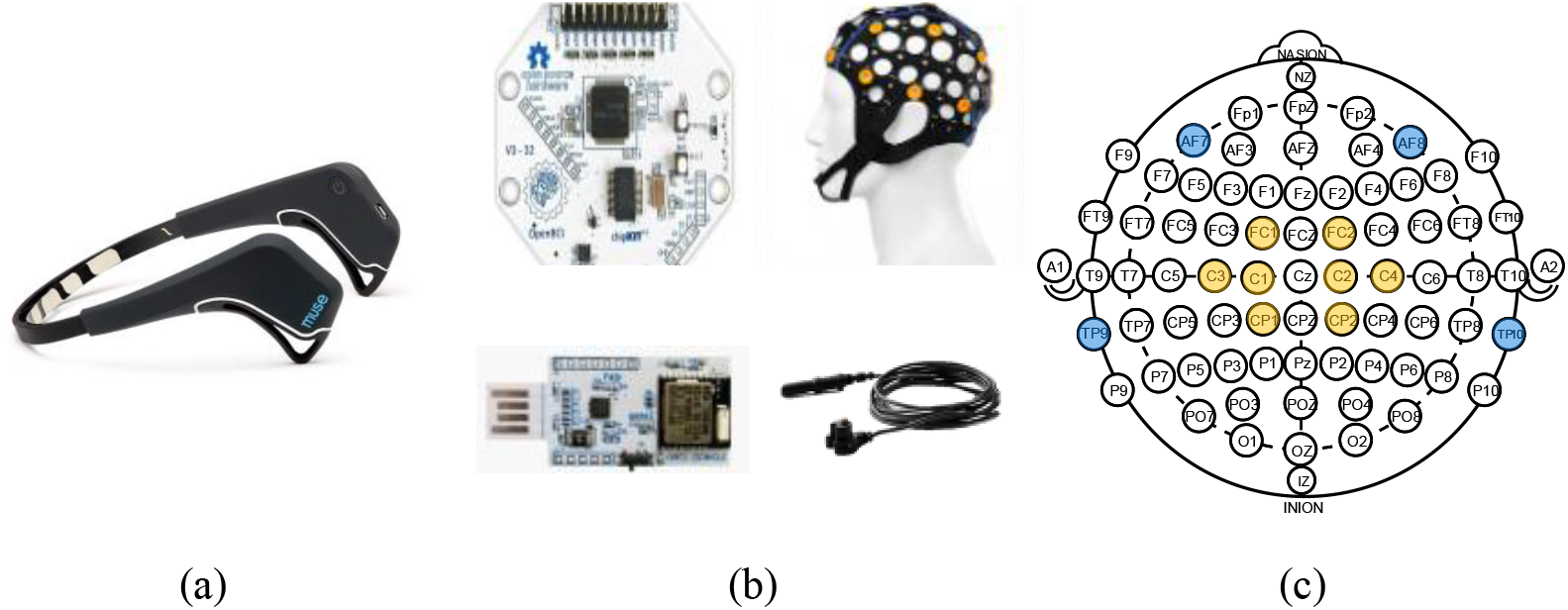
Devices and their channel configurations. (a) The Muse EEG headband. (b) The Cyton biosensing 8-channel bio-amplifier (top-left), including its USB dongle (bottom-left), MCScap 10-20 marked textile cap (top-right), and MCScap-E electrode (bottom-right). (c) Channel configuration of both EEG devices, based on the modified expanded 10-20 system, where blue markers indicate Muse and Yellow markers indicate OpenBCI electrode locations.

### 2.2. Participants and Experimental Procedure

The study recruited six healthy participants (five right-handed, age: 27.8 ± 12.2). Each participant completed two independent recording sessions with each of the Muse headband and OpenBCI system. These sessions were conducted in a noise-free room to minimize external disturbances. Participants were asked to sit comfortably in a chair with their arms resting at their sides. Each recording session consisted of six test sessions, each of which comprised 40 sequential trials. “Table 1” presents the motor tasks performed during the experiment, detailing the tasks names and corresponding participant actions.

**Table 1.** Motor tasks descriptions.

| Motor task | Participant action |
| --- | --- |
| Left knee motor imagery (LKMI) | Imagining the extension of the left knee |
| Right knee motor imagery (RKMI) | Imagining the extension of the right knee |
| Left knee motor execution (LKME) | Performing a slow extension of the left knee |
| Right knee motor execution (RKME) | Performing a slow extension of the right knee |
| Left ankle motor imagery (LAMI) | Imagining the dorsiflexion of the left ankle |
| Right ankle motor imagery (RAMI) | Imagining the dorsiflexion of the right ankle |

“Figure 3” illustrates the trial structure. Each trial began with a cue phase, where a 0.15s beep signaled the trial’s start, followed by a 5-second visual stimulus (“CUE” with a red circle on a LED display), allowing participants to prepare and adjust. Next, the rest/motor task phase started with another 0.15s beep, followed by a randomly assigned 3.3s, 3.8s, or 4.3s stimulus displaying either “REST” (yellow circle) or a motor task (green circle). ME tasks were excluded in this phase to allow participants to reset their knee position before the next movement. Another motor/rest task phase followed, mirroring the previous phase but including ME tasks. EEG epochs were extracted from phases 2 and 3, excluding the first 0.2s and last 0.1s to reduce transition effects, resulting in 3s, 3.5s, or 4s epochs. Finally, the post-task cue phase, identical to initial cue, provided a brief rest before the next trial.

**Figure 3.**
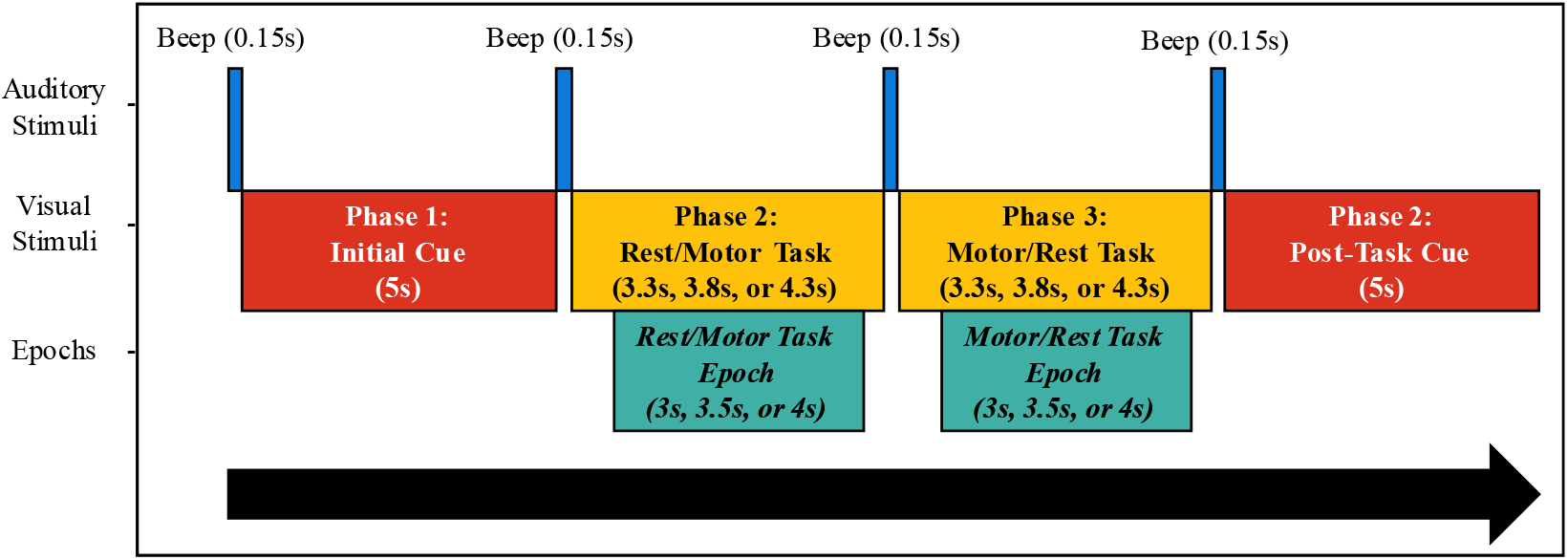
Schematic representation of the trial structure, detailing the sequence of auditory and visual stimuli across different phases. The trial begins with a preparatory cue, followed by alternating task and rest phases, and concludes with a post-task cue. EEG epochs were extracted by excluding the first 0.2s after the onset of the visual stimulus and the last 0.1s before its offset.

### 2.3. Preprocessing and Segmentation

To ensure uniform temporal resolution across datasets, the recordings of the two EEG systems were resampled to a common sampling rate of 256 Hz using polyphase rational resampling [16]. Then, a 4th-order Butterworth bandpass filter was applied in the frequency range of 8-30 Hz, covering the alpha (8-13 Hz) and beta (14-30) frequency bands, which are strongly associated with motor tasks [17]. The 4th-order Butterworth filter was chosen for its maximally flat (ripple-free) passband and balanced amplitude selectivity, which helps minimize distortion within the passband [18].

Over the course of six test sessions, each participant completed a total 240 epochs per motor task and 1440 epochs for the rest task per recording device. Each epoch was further segmented into one-second windows (256 samples) with a 0.75s overlap (192 samples), improving temporal resolution, addressing EEG non-stationarity, and increasing training samples per class. “Table 2” details the epoch and segment distribution per participant and device.

**Table 2.**
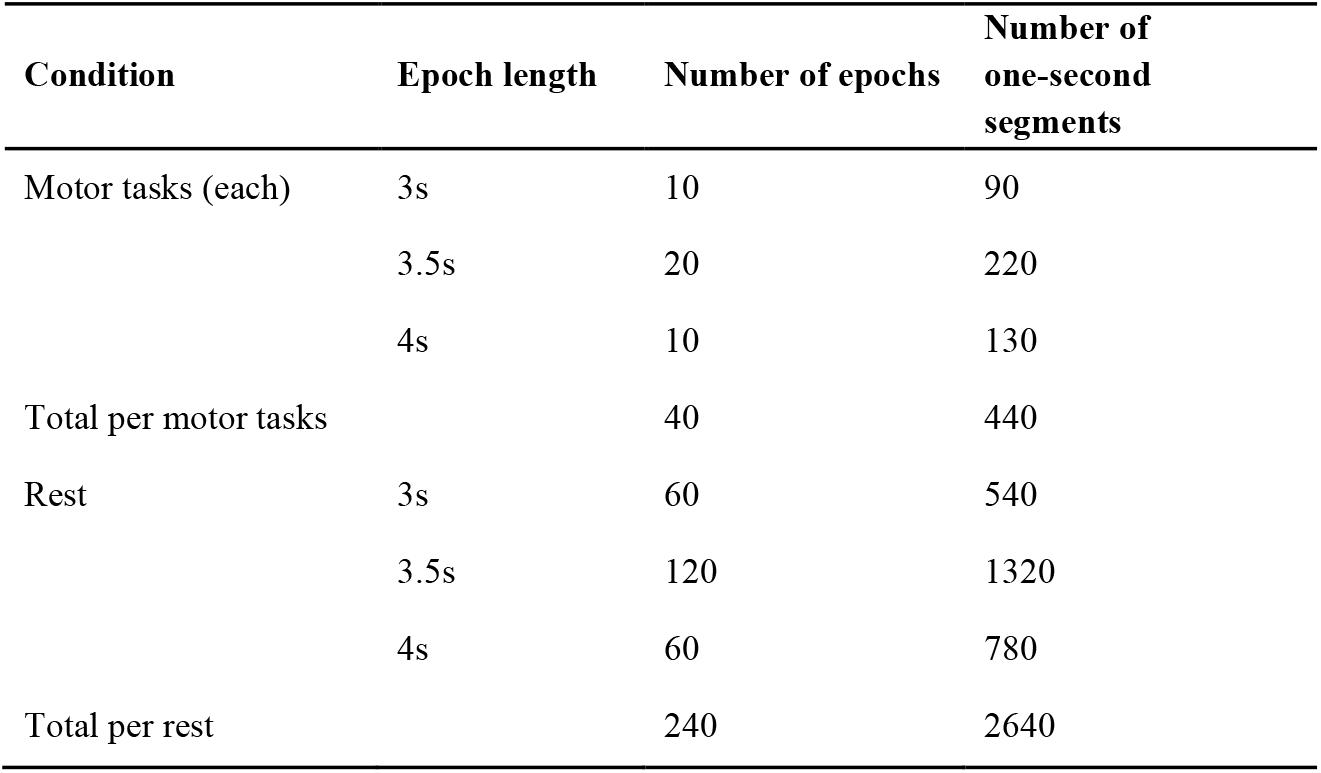
Epoch and segment distribution per subject, per device.

| **Condition** | **Epoch length** | **Number of epochs** | **Number of one-second segments** |
| --- | --- | --- | --- |
| Motor tasks (each) | 3s | 10 | 90 |
|  | 3.5s | 20 | 220 |
|  | 4s | 10 | 130 |
| Total per motor tasks |  | 40 | 440 |
| Rest | 3s | 60 | 540 |
|  | 3.5s | 120 | 1320 |
|  | 4s | 60 | 780 |
| Total per rest |  | 240 | 2640 |

### 2.4. Feature Extraction

Feature extraction was conducted in three primary domains of time (TD), frequency (FD), and time-frequency (TFD), leveraging established techniques from prior research to ensure comprehensive signal representation. A complete list of the extracted features, categorized by their respective approaches is provided in “Table 3”. Detailed description of the features is given in the following.

**Table 3.**
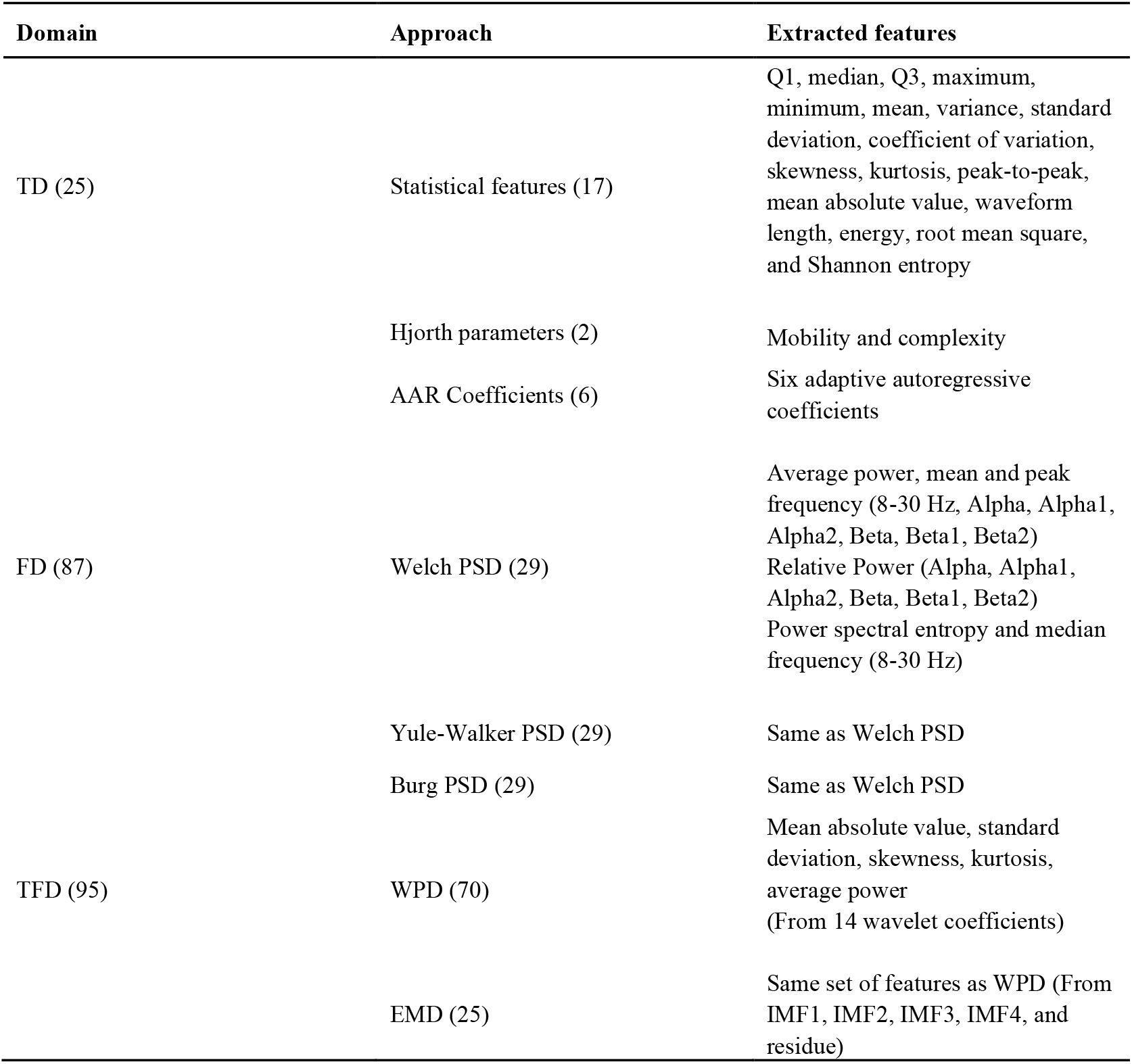
Extracted features categorized by domain and method.

| Domain | Approach | Extracted features |
| --- | --- | --- |
| TD (25) | Statistical features (17) | Q1, median, Q3, maximum, minimum, mean, variance, standard deviation, coefficient of variation, skewness, kurtosis, peak-to-peak, mean absolute value, waveform length, energy, root mean square, and Shannon entropy |
|  | Hjorth parameters (2) | Mobility and complexity |
|  | AAR Coefficients (6) | Six adaptive autoregressive coefficients |
| FD (87) | Welch PSD (29) | Average power, mean and peak frequency (8-30 Hz, Alpha, Alpha1, Alpha2, Beta, Beta1, Beta2)<br>Relative Power (Alpha, Alpha1, Alpha2, Beta, Beta1, Beta2)<br>Power spectral entropy and median frequency (8-30 Hz) |
|  | Yule-Walker PSD (29) | Same as Welch PSD |
|  | Burg PSD (29) | Same as Welch PSD |
| TFD (95) | WPD (70) | Mean absolute value, standard deviation, skewness, kurtosis, average power<br>(From 14 wavelet coefficients) |
|  | EMD (25) | Same set of features as WPD (From IMF1, IMF2, IMF3, IMF4, and residue) |

#### 2.4.1. Time Domain Features

The extracted TD features were categorized into three groups of statistical features, Hjorth parameters, and adaptive autoregressive (AAR) coefficients. We denote *x*[*n*] as the EEG signal with samples *x*_1_ to *x*_*N*_, where *N* = 256.

Statistical features [19, 20, 21] included first quartile (Q1), median, third quartile (Q3), maximum, minimum, and peak-to-peak amplitude values. Additionally, we computed mean (*μ*), variance (*σ*^2^), standard deviation (*σ*), coefficient of variation (*σ/μ*), skewness, and kurtosis to quantify signal dispersion and shape. Other extracted features included mean absolute value, waveform length, energy, root mean square, and Shannon entropy [22].

Hjorth mobility and complexity were extracted to describe signal dynamics [23]. Let the first-order difference of *x*[*n*] be:

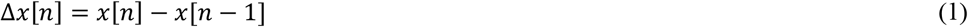

Then, Hjorth mobility would be expressed as:

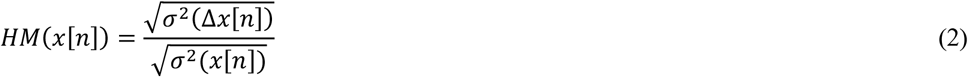

And Hjorth complexity is:

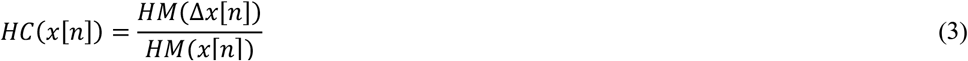

The AAR model represents an EEG segment as a sum of weighted past values with time-varying coefficients expressed as:

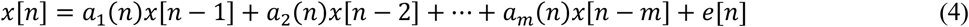

Where *a*_*i*_(*n*) are time-dependent autoregressive (AR) coefficients, *m* is the model order, and *e*[*n*] is the residual error. We used a model order of six, as higher orders did not significantly enhance classification performance [24, 25]. The extracted features included six coefficients, *a*_1_(*n*) to *a*_6_(*n*), to capture EEG signal temporal dependencies.

#### 2.4.2. Frequency Domain Features

The power spectral density (PSD) was estimated using Welch’s non-parametric method [26] and two parametric approaches: the Yule-Walker and Burg AR models [27]. Features were extracted from all three methods for comparison. Welch’s method involves segmenting EEG signals into overlapping windows, applying a Hamming window, and averaging periodograms. Here, the signals were segmented into three 128-sample windows with 50% overlap, and PSD values were normalized by total power. The Yule-Walker and Burg methods both estimate the PSD by modelling EEG signals as AR processes of order *P*, where the spectral content is shaped by AR coefficients and noise variance. In the Yule-Walker method, the PSD is derived from the AR model’s frequency response as:

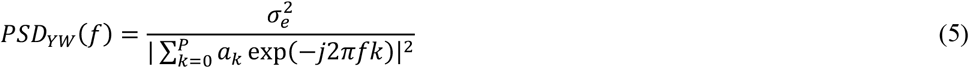

Where 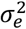 is the noise variance and *a*_*k*_ are the AR coefficients. The Burg method, instead, iteratively estimates reflection coefficients to minimize forward and backward prediction errors. To determine the optimal AR model order *P* for both methods, the Akaike Information Criterion (AIC) [28] was applied, selecting the order that minimizes:

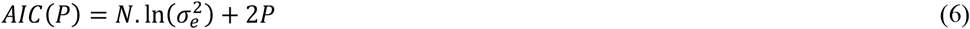

Where, *N* = 256 is the number of EEG samples in each segment, and 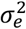 is the estimated noise variance.

Features were extracted from PSDs across EEG frequencies (8–30 Hz), including alpha and beta bands associated with motor control [17, 29]. To capture finer spectral variations relevant to the motor task classification, sub-bands divisions within alpha and beta were also considered: alpha1 (8-10.5 Hz), alpha2 (10.5-13 Hz), beta1 (14-22 Hz), and beta2 (22-30 Hz). The extracted features included average power, mean, median, and peak frequency within the predefined bands [19]; relative power of all sub-bands normalized to the total power in 8-30 Hz [30], and power spectral entropy within the full range of 8-30 Hz [31].

#### 2.4.3. Time-Frequency Domain Features

Wavelet packet decomposition (WPD) and Empirical Mode Decomposition (EMD) were used for TFD analysis. WPD extends the Discrete Wavelet Transform (DWT) [32] by decomposing both low and high frequency components at each level. At each decomposition level, the signal is passed through high-pass and low-pass filters, then down sampled by a factor of two. Unlike DWT, which produces *j* + 1 sets of wavelet coefficients at level *j*, WPD generates 2^*j*^ sets, offering improved frequency resolution [33]. EMD decomposes the signal into intrinsic mode functions (IMFs) through an iterative sifting process [34]. The highest frequency component (IMF1) is extracted first, followed by progressively lower-frequency components, leaving a residual signal at the final stage. For each EEG segment, we performed a three-level WPD decomposition using the Sym4 wavelet function, as it produced the best overall performance [33], yielding eight wavelet coefficient sets. From each set, we extracted mean absolute value, standard deviation, skewness, kurtosis, and average power [33, 35]. Similarly, for EMD, we selected the first four IMFs and the final residue, extracting the same statistical features as in WPD.

By applying diverse feature extraction methods, we aimed to assess their relevance in motor tasks classification. To ensure a robust feature space, all features were extracted per EEG segment from each channel. With 207 features per segments, the total feature count per one-second segments was 149 × 4 for the low-channel device (Muse) and 149 × 8 for the higher-channel device (OpenBCI).

### 2.5. Feature Selection and Dimension Reduction

To prevent overfitting and reduce computational complexity, feature selection was performed using the minimum redundancy maximum relevance (mRMR) algorithm [36], based on mutual information [36] to remove irrelevant or redundant features. Mutual information quantifies how knowledge of one variable reduces uncertainty about another. Given two random variables *A* and *B*, their mutual information is defined as:

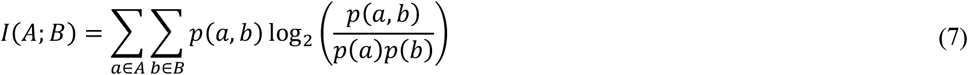

Where *p*(*a, b*) is the joint probability of *A* = *a* and *B* = *b*, and *p*(*a*) and *p*(*b*) are the marginal probabilities of *A* = *a* and *B* = *b*, respectively. Given a dataset *X* ∈ ℝ^*M*×*N*^ (with *M* samples and *N* features per sample) and a label vector *Y* = {*y*_1_, …, *y*_*M*_}, where *y*_*i*_ ∈ {*C*_1_, *C*_2_} represents the class label, the relevance of a feature *X*_*i*_ with respect to the class label *Y* is defined as *I*(*X*_*i*_; *Y*), and the redundancy between two features *X*_*i*_ and *X*_*j*_ is defined as *I*(*X*_*i*_; *X*_*j*_). The mRMR criterion selects a subset of features *S* that maximizes the relevance to the class label and minimizes redundancy among selected features. A greedy algorithm first selects the feature with the highest relevance, then iteratively adds features by optimizing:

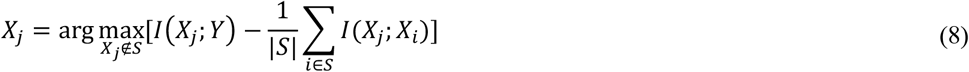

This continues until the desired number of features is selected. Using mRMR, 100 features were finally selected for each device across all channels. After feature selection, Principal Component Analysis (PCA) [37] was applied to further reduce dimensionality while preserving variance. PCA transforms the selected feature matrix X into a new space:

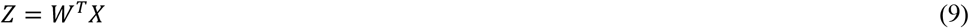

Where *W* is the matrix of eigenvectors of the covariance matrix of *X*, and *Z* represents the transformed features. The top components capturing at least 95% of cumulative variance were retained, balancing dimensionality reduction with information preservation.

### 2.6. Classification

A subject-specific approach was used for classification, where binary classifiers were trained separately for each participant. As shown in “Table 4”, classification was performed across nine scenarios, categorized into three groups. We used five widely adopted classifiers: Support Vector Machine (SVM), Linear Discriminant Analysis (LDA), k-Nearest Neighbours (KNN), Random Forest (RF), and Adaptive Boosting (AdaBoost). For model training, the dataset was randomly split into 10 folds of identical sizes. Seven folds were used to train the initial classifier, two folds for hyperparameter tuning, and one final fold for evaluation. This process was repeated 100 times, and the final classification accuracy was found as the mean performance across all runs.

**Table 4.**
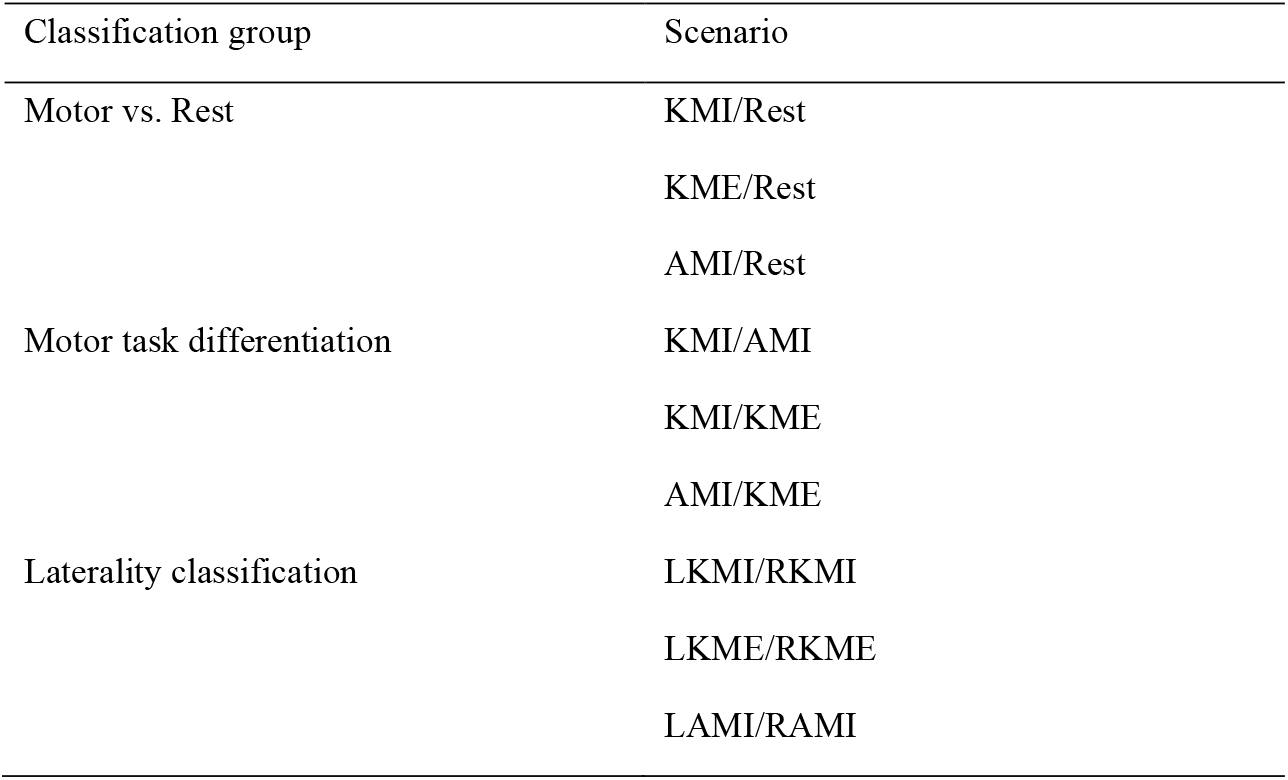
Binary classification scenarios.

| Classification group | Scenario |
| --- | --- |
| Motor vs. Rest | KMI/Rest |
|  | KME/Rest |
|  | AMI/Rest |
| Motor task differentiation | KMI/AMI |
|  | KMI/KME |
|  | AMI/KME |
| Laterality classification | LKMI/RKMI |
|  | LKME/RKME |
|  | LAMI/RAMI |

SVM constructs an optimal hyperplane to maximize class separation, with hyperparameter tuning performed for kernel type and regularization [38]. LDA assumes Gaussian-distributed classes with a shared covariance matrix and determines a linear decision boundary, with shrinkage parameters optimized during training [39]. KNN assigns labels based on the majority vote of *k*- nearest neighbors, where *k* and the distance metric are selected through cross-validation [40]. RF is an ensemble classifier that aggregates multiple decision trees, with the number of trees and feature selection criteria tuned to enhance generalization [41]. AdaBoost builds an additive ensemble of weak RFs trained sequentially, then after each stage, misclassified samples receive higher weights so the next weak RF focuses on hard cases. Final predictions are a weighted vote across weak RFs, with stage weights tied to their training loss (learning rate controls shrinkage). We tuned boosting rounds, learning rate, and the internal capacity of each weak RF, using cross-validation/early stopping to prevent overfitting [42].

## 3. Results

### 3.1. Feature Selection Outcomes

“Figure 4” depicts the overall distribution of the selected features across all participants and classification tasks. The inner ring of each pie chart shows the proportion of features from different feature domains, while the outer ring displays the distribution across the extraction approaches within each domain. The left pie chart corresponds to the Muse device, and the right pie chart represents the OpenBCI device.

**Figure 4.**
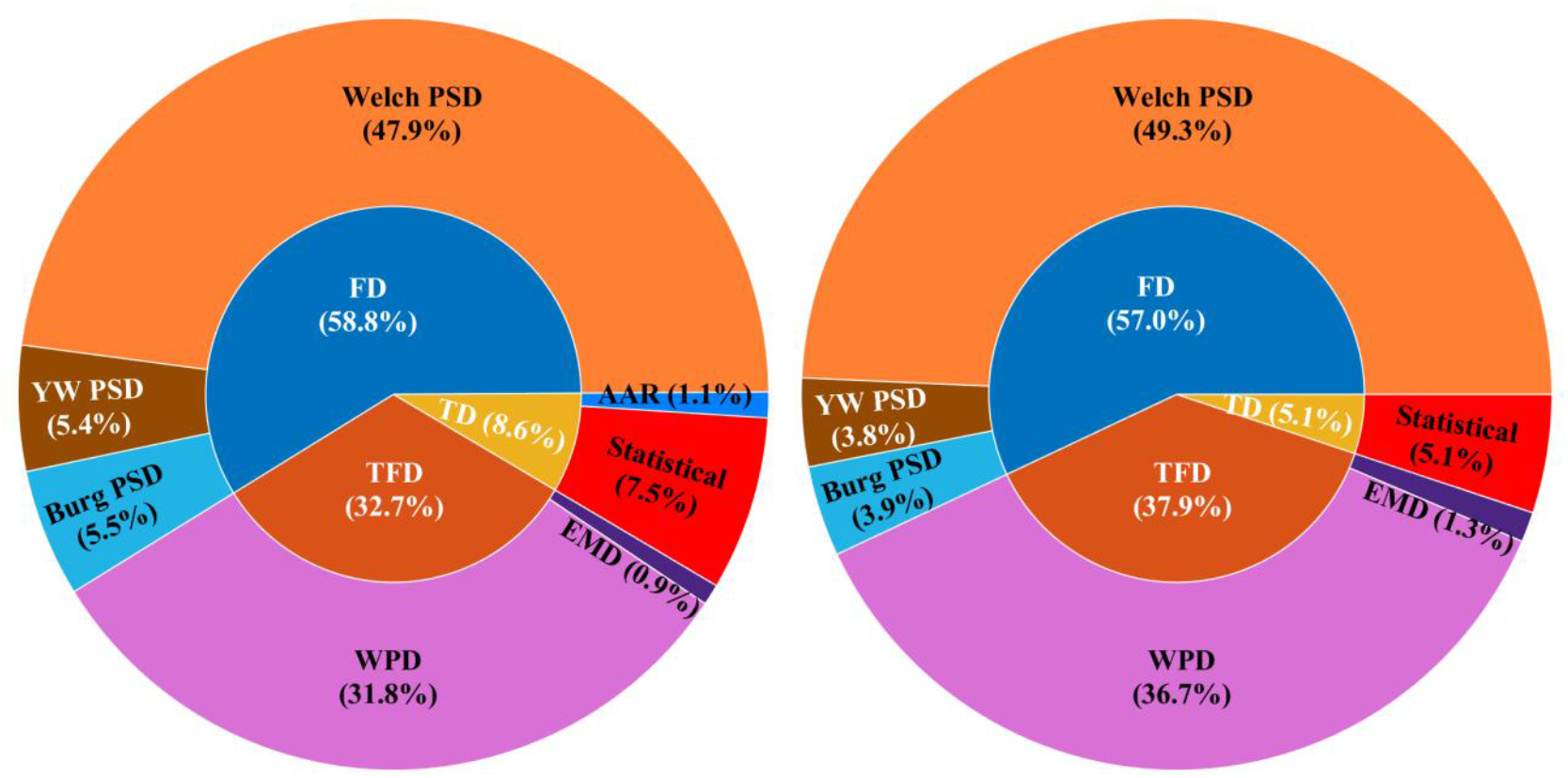
Distribution of the selected features across feature domains (innter ring) and approaches (outer ring) for both low-channel (Muse) and higher-channel (OpenBCI) devices.

### 3.2. Mutual Information Analysis by Channel

To investigate the significance of the selected features, we computed their mutual information scores for three classification groups: motor vs. rest, motor task differentiation, and laterality. These scores were aggregated across participants and scenarios, then summarized per channel, highlighting how each channel and feature extraction approach contributed to the total mutual information. “Figure 5” illustrates these results via stacked column charts for the Muse device (a) and the OpenBCI system (b).

**Figure 5.**
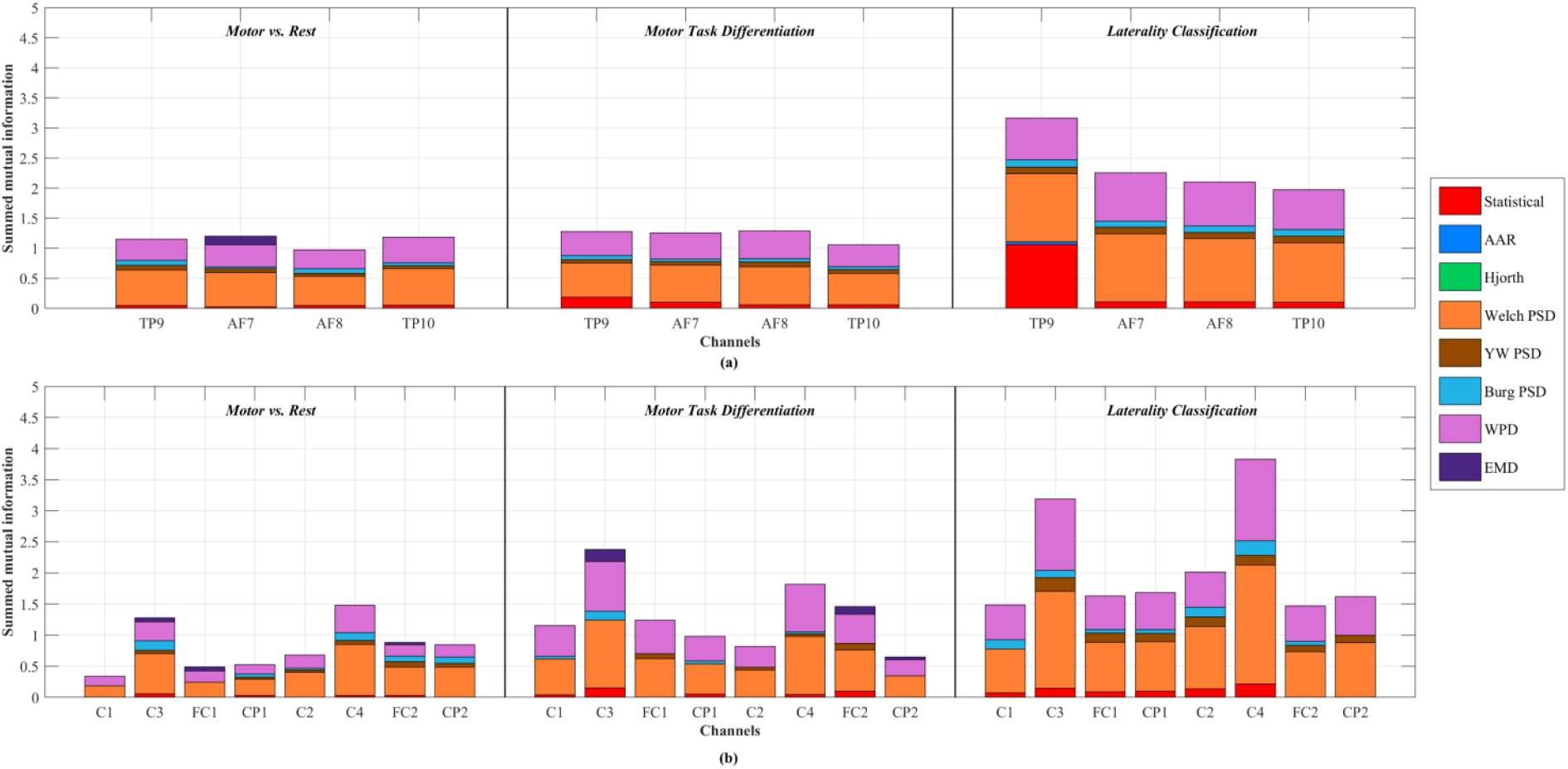
Stacked column plots of summed mutual information per channel and approach. (a) The Muse (low-channel) device, and (b) the OpenBCI (higher-channel) device.

### 3.3. Classification Performance

Each classification scenario was repeated for 100 iterations to account for variability, and the final accuracy was obtained by averaging confusion matrices over all iterations and participants. “Figure 6” presents a heatmap of the average classification accuracies for each of the nine scenarios using the Muse device. “Figure 7” shows the similar heatmap for the OpenBCI device.

**Figure 6.**
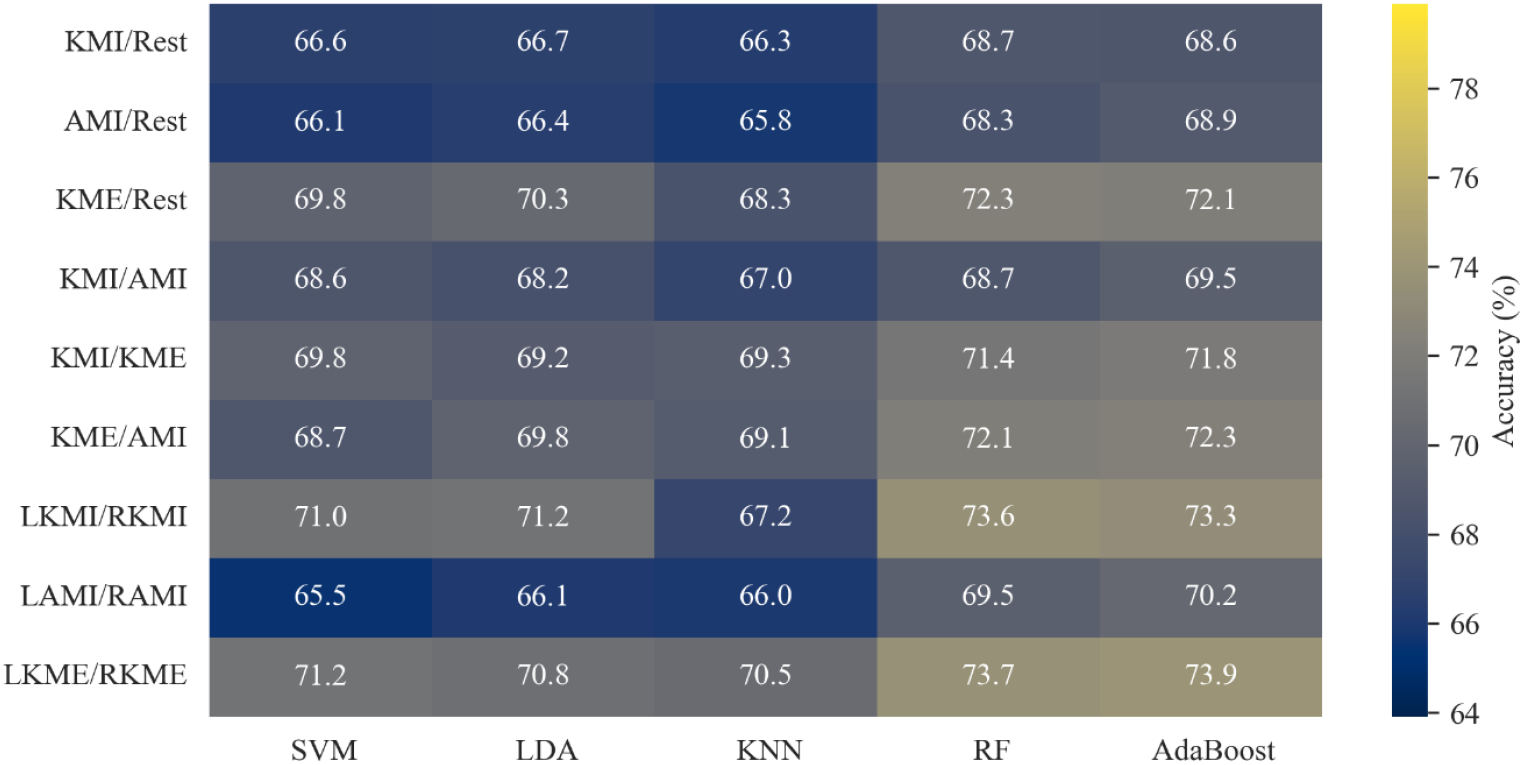
Heatmap of mean accuracies for each of the nine scenarios (rows) and five classifiers (columns) using the Muse (low-channel) device.

**Figure 7.**
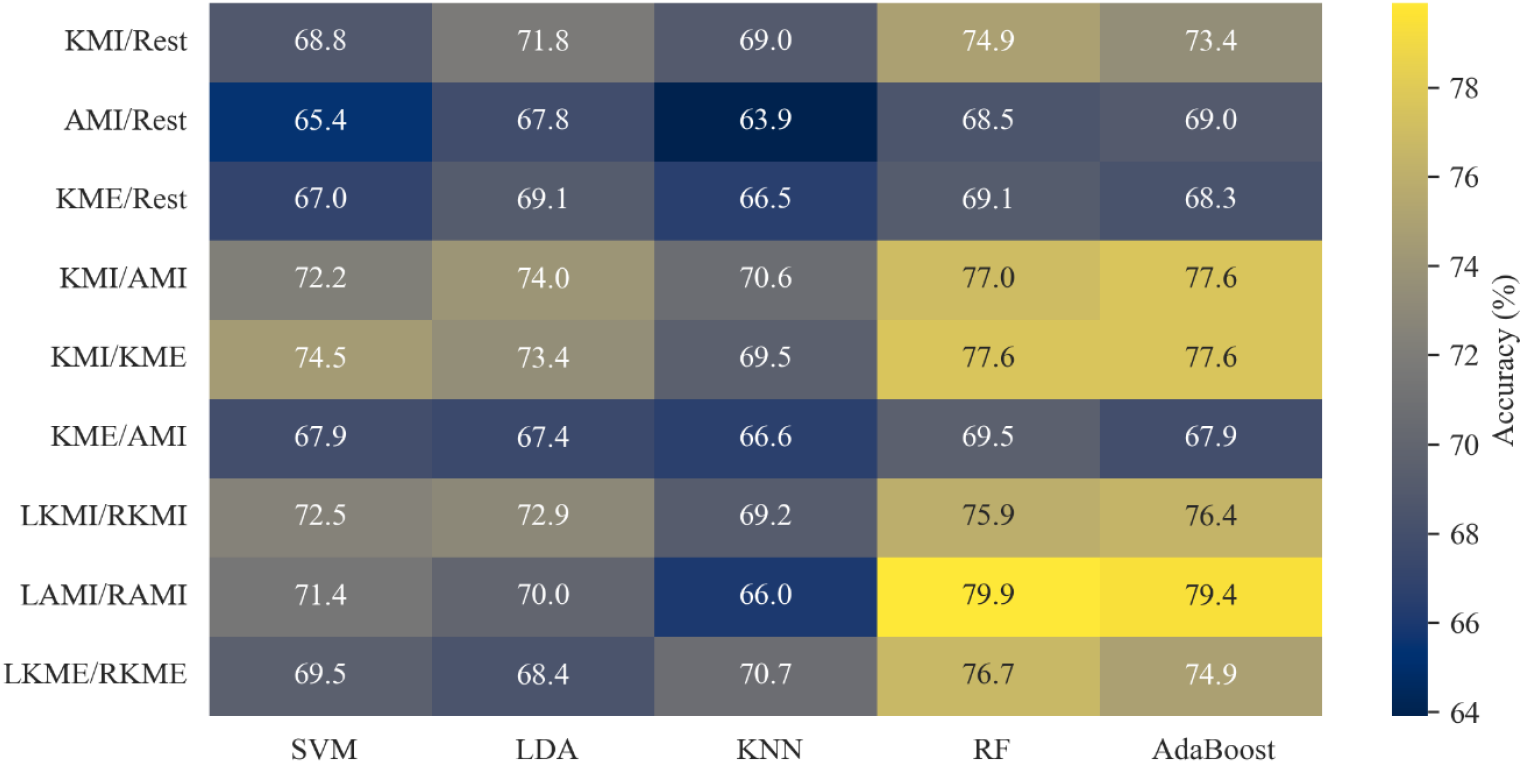
Heatmap of mean accuracies for each scenario and classifier with the OpenBCI (higher-channel) device.

### 3.4. Comparison of the two devices

“Figure 8” shows the boxplots of accuracies for each of the five classifiers, averaged over all task scenarios, for both devices. As expected, the OpenBCI device has superior performance across all classifiers. The relative superiority of the OpenBCI system over Muse is written for each classifier as a percentage alongside the boxplot. “Figure 9” provides a similar set of boxplots for the nine task scenarios (averaged over all classifiers), as well as the corresponding relative superiority of the OpenBCI system.

**Figure 8.**
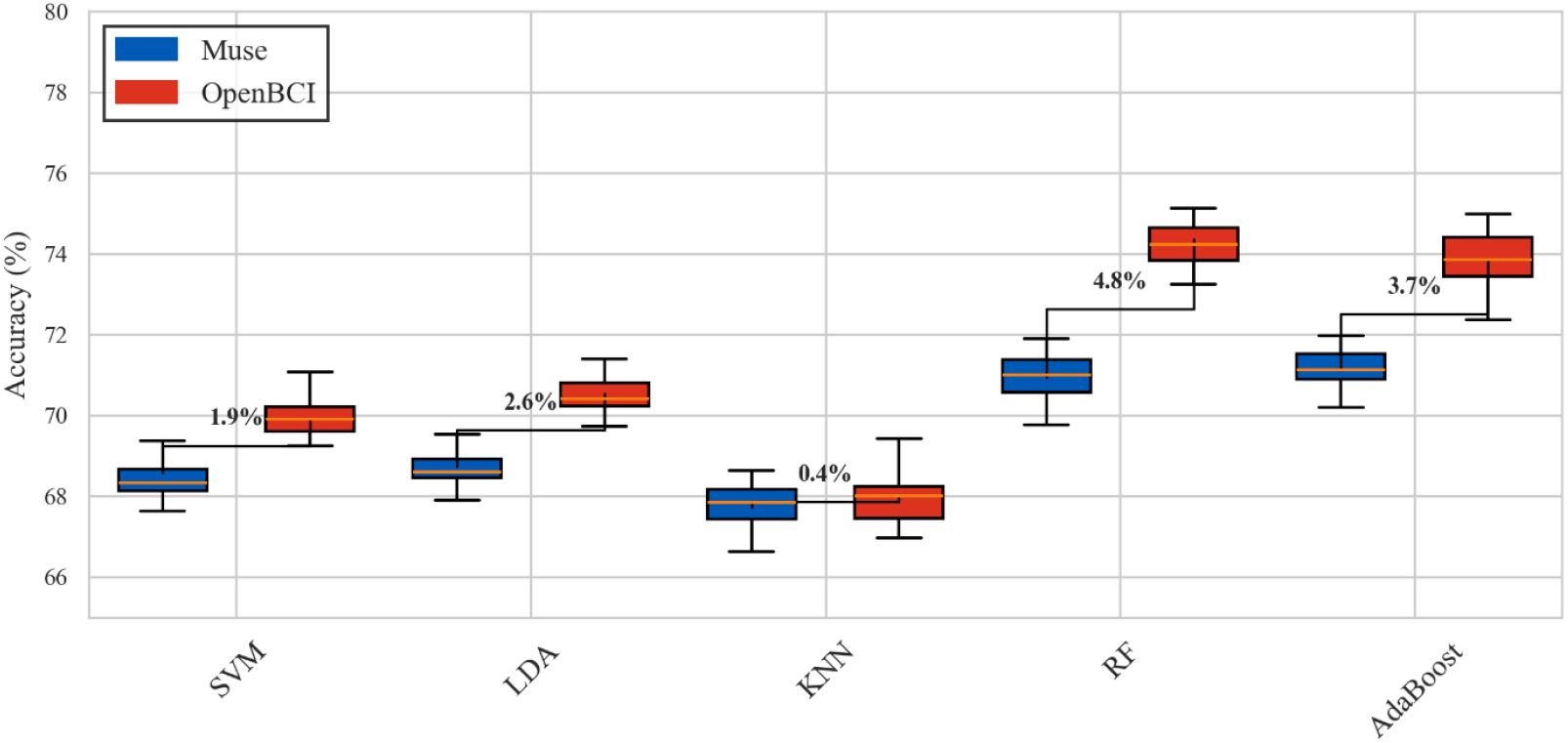
Boxplots of classification accuracies for the five classifiers (both devices), averaged over task scenarios, with the relative superiority of the OpenBCI device over the Muse system.

**Figure 9.**
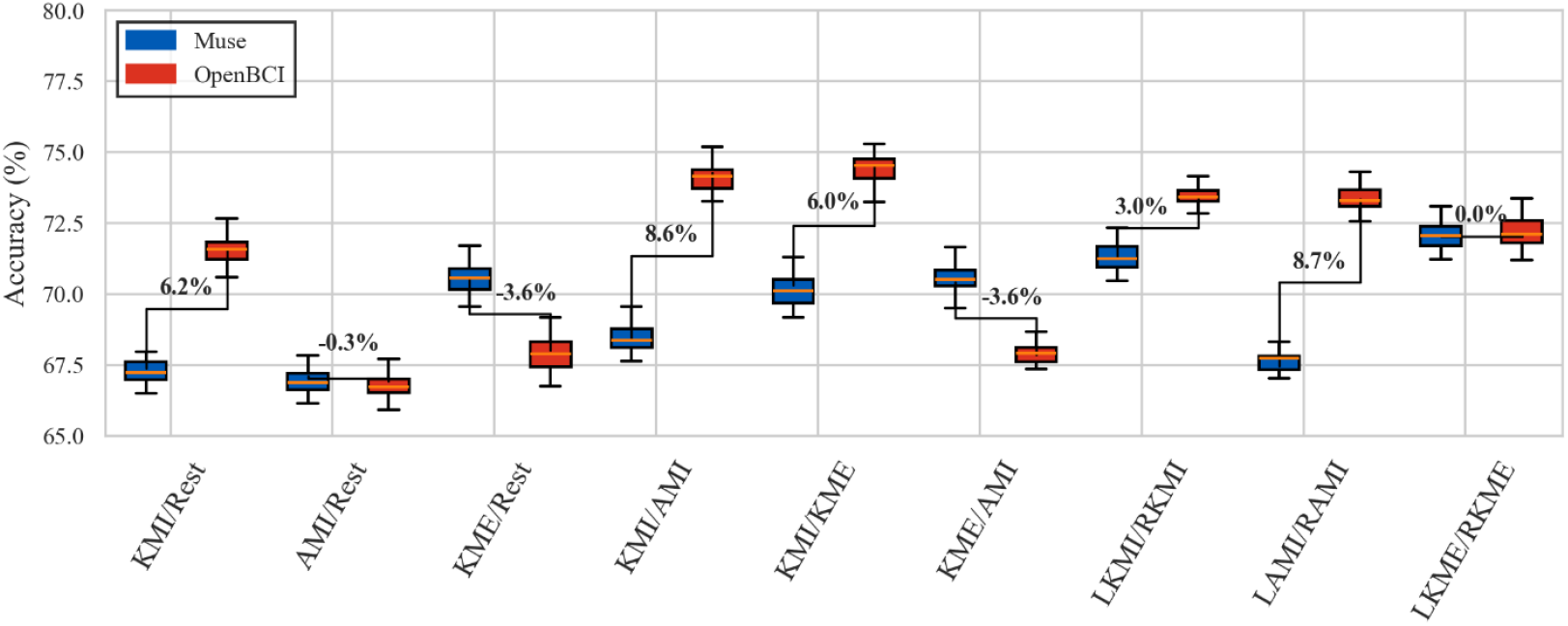
Boxplots of the classification accuracies of the nine task scenarios (both devices), averaged over all classifiers, as well as the telative superiority of the OpenBCI system.

## 4. Discussion

The feature selection analysis, shown in “Figure 4”, revealed that FD features dominate the classification of motor related tasks, followed by TFD features, while TD features contribute the least. This aligns with previous studies highlighting the critical role of oscillatory dynamics in motor related EEG activity, particularly in the 8–30 Hz range. Variations in power within this range are key for distinguishing MI tasks, confirming that spectral features are generally more informative than TD features [43, 44, 45]. Further analysis indicates that signal differences in MI primarily occur in frequency-specific dynamics, which time-domain features fail to capture. Despite integrating temporal information, TFD features still contribute less than FD features, emphasizing that frequency-specific changes, rather than simple signal amplitude fluctuations, are crucial for accurate MI classification.

Among the FD approaches, Welch’s PSD emerged as the most frequently selected approach, underscoring the effectiveness of non-parametric spectral estimations in capturing transient motor-related EEG changes. In contrast, parametric methods such as Yule-Walker and Burg contributed less, which highlights the advantage of non-parametric methods in real-world applications where noise can obscure signal features [27]. Moreover, the pattern of selected features showed only slight variations between the two devices studied. No significant differences in feature selection outcomes were observed between the Muse and OpenBCI devices, suggesting that feature selection performance was not substantially affected by the device’s channel count.

The mutual information analysis by channel, shown in “Figure 5” reveals a relatively clear pattern in the contribution of individual channels across classification tasks or the Muse device, all four channels contributed relatively uniformly, except in the laterality classification task, where the TP9 channel was notably more significant. This increased significance may be attributed to the fact that the subjects in this study were predominantly right-handed. In contrast, the OpenBCI system consistently highlighted the C3 and C4 channels as most significant across all tasks, reinforcing the known importance of these central motor cortical electrodes in MI classification [45, 46]. Laterality classification has the highest summed mutual information, reinforcing that the features provide the most useful information for distinguishing left/right motor-related tasks. In terms of feature group contributions, the share of feature types across channels appears to be almost uniform, suggesting that the significance of the feature type is not strongly dependent on the specific channel. Welch PSD was consistently the most significant feature group, followed by time-frequency WPD features, corroborating results from “Figure 4”. The only notable exception was observed for the TP9 channel of the Muse device during laterality classification, where TD statistical features were more important than FD and TFD features. This suggests that temporal differences in this channel were particularly relevant for distinguishing between left and right motor tasks, which may reflect unique neural dynamics captured by the Muse device in this specific classification context.

The classification performance results, shown in “Figure 6” for the Muse device and “Figure 7” for the OpenBCI system, indicate that both devices yield a similar pattern of mean accuracy across the binary tasks, such as motor/rest, motor task differentiation, and laterality detection. The mean accuracy for these tasks is around 70%, consistent with previously reported results for MI classification using EEG signals [7,8,]. This suggests that both systems providing comparable results. Notably, both devices perform slightly better with RF and AdaBoost classifiers, likely because these ensemble methods handle variability and noise in EEG data more effectively by reducing overfitting and improving generalization. There are no significant differences in accuracy between task types, and no strong performance advantage is observed between devices at this stage of analysis.

## 5. Conclusions

The comparative analysis of the two devices, as illustrated in “Figure 8” and “Figure 9”, reveals relatively close performance, with the OpenBCI system demonstrating a slight overall advantage. This superiority is most evident when using ensemble-based classifiers such as RF and AdaBoost, which may benefit from OpenBCI’s higher spatial resolution. In contrast, other classifiers show minimal gains, suggesting that the added spatial information does not universally enhance classification accuracy. Task-specific comparisons further support this observation, with differences in accuracy between the two devices ranging from -3.6% to 8.6% across various scenarios. As a concluding remark, the results indicate that for binary classification of lower limb motor tasks, the more affordable Muse device can perform comparably to the lab-scale OpenBCI system, making it a viable option for portable and cost-sensitive BCI applications.

## Abbreviations

AAR: Adaptive Autoregressive
AR: Autoregressive
BCI: Brain Computer Interfaces
CNS: Central Nervous System
DWT: Discrete Wavelet Transform
EEG: Electroencephalography
EMD: Empirical Mode Decomposition
FD: Frequency Domain
IMF: Intrinsic Mode Function
KNN: K-Nearest Neighbours
LAMI: Left Ankle Motor Imagery
LDA: Linear Discriminant Analysis
LKME: Left Knee Motor Execution
LKMI: Left Knee Motor Imagery
ME: Motor Execution
MI: Motor Imagery
mRMR: Minimum Redundancy Maximum Relevance
PCA: Principal Component Analysis
PSD: Power Spectral Density
RAMI: Right Ankle Motor Imagery
RF: Random Forest
RKME: Right Knee Motor Execution
RKMI: Right Knee Motor Imagery
SVM: Support Vector Machine
TD: Time Domain
TFD: Time-Frequency Domain
WPD: Wavelet Packet Decomposition

## 7. Declarations

## Ethics approval and consent to participate

The experimental protocol was approved by the ethics committee of Iran University of Medical Sciences (IR.IUMS.REC.1403.455) in accordance with the Declaration of Helsinki. All participants provided informed consent.

## Competing Interests

The authors declare no competing interests.

## Funding

Not applicable.

## Authors’ contributions

– Parsa Bahramsari: Methodology, Software, Investigation, Formal analysis, Writing original draft.
– Saeed Behzadipour: Conceptualization, Methodology, Supervision, Writing-Review & Editing.

## Availability of data and materials

The data described in this Data note can be freely and openly accessed on Zenodo under https://doi.org/10.5281/zenodo.16455324

## Competing interests

The authors declare that they have no competing interests.

## Acknowledgements

We would like to thank all participants of our experiments and the members of the Djawad Mowafaghian Research Center in Neuro-Rehabilitation Technologies.

## Notes

### Competing Interest Statement

The authors have declared no competing interest.

